# 3D Flexible Refinement: Structure and Motion of Flexible Proteins from Cryo-EM

**DOI:** 10.1101/2021.04.22.440893

**Authors:** Ali Punjani, David J. Fleet

## Abstract

Single particle cryo-EM excels in determining static structures of biological macromolecules such as proteins. However, many proteins are dynamic, with their motion inherently linked to their function. Recovering the continuous motion and detailed 3D structure of flexible proteins from cryo-EM data has remained an open challenge. We introduce *3D Flexible Refinement* (3DFlex), a motion-based deep neural network model of continuous heterogeneity. 3DFlex directly exploits the knowledge that conformational variability of a protein is often the result of physical processes that transport density over space and tend to conserve mass and preserve local geometry. From 2D image data, the 3DFlex model jointly learns a single canonical 3D map, latent coordinate vectors that specify positions on the protein’s conformational landscape, and a flow generator that, given a latent position as input, outputs a 3D deformation field. This deformation field convects the canonical map into appropriate conformations to explain experimental images. Applied to experimental data, 3DFlex learns non-rigid motion spanning several orders of magnitude while preserving high-resolution details of secondary structure elements. Further, 3DFlex resolves canonical maps that are improved relative to conventional refinement methods because particle images contribute to the maps coherently regardless of the conformation of the protein in the image. Together, the ability to obtain insight into motion in macromolecules, as well as the ability to resolve features that are usually lost in cryo-EM of flexible specimens, will provide new insight and allow new avenues of investigation into biomolecular structure and function.

## 1 Introduction

Proteins form the molecular machinery of the cell. They are inherently dynamic, often exhibiting a continuous landscape of conformations, with motion tightly coupled to function. Methods that uncover protein motion and the conformational landscape have the potential to illuminate fundamental questions in structural biology, and to enhance the ability to design therapeutic molecules that elicit specific functional changes in a target protein.

Single particle cryo-EM collects thousands of static 2D protein particle images that, in aggregate, may span the target protein’s 3D conformational space. Cryo-EM therefore holds great promise for uncovering both the atomic-resolution structure and motion of biologically functional moving parts [19]. Nevertheless, methods for resolving continuous motion and structure from static 2D images have remained elusive.

Established high-resolution cryo-EM refinement methods assume rigidity of the target molecule, and result in blurred, low-resolution density for flexible regions [12, 29, 35]. Methods with spatially adaptive regularization [30, 31] mitigate the adverse effects of local motion on rigid refinement, but do not estimate the motion *per se*. Local refinement and multi-body methods [23] use masks to focus on a sub-region of a protein, but only provide improved resolution on rigid parts of relatively high molecular weight [37]. Subspace methods [28, 42] approximate a particle’s space of conformations only as a linear combination of basis density maps, and without an underlying model of motion.

The development of a method that can uncover protein motion in the presence of continuous flexibility and thereby improve the resolution of fine structural details has been a long standing open problem, but effective solutions face several key challenges. First, there are a large number of unknowns that must be jointly estimated from the data, including the 3D structure of the density map, a representation of the space of conformational changes, and the position of each particle image on that conformational landscape. Second, the protein motion and the conformational landscape are generally non-linear. Third, in order to resolve 3D map details beyond what conventional (static) reconstructions can provide, structural information must be aggregated from many different conformations. Finally, despite the high levels of noise, and computational difficulty of the underlying optimization problem, the unknowns must be estimated with sufficient precision to enable recovery of high-resolution details.

We introduce *3D Flexible Refinement* (3DFlex), a deep neural network model of continuously flexible protein molecules. 3DFlex is a motion-based heterogeneity model that directly exploits the knowledge that most conformational variability of a protein is a result of physical processes that transport density over space and tend to conserve mass and preserve the local geometry of the underlying structure. This is in contrast to density-based techniques that model conformational variation as a manifold of 3D density maps without a physically plausible motion model [9, 28, 42, 43, 44]. Some recently proposed techniques have sought to model motion, albeit in limited forms [3, 17]. The formulation of 3DFlex is based on a generative architecture that captures conformational variability in terms of a single high-resolution “canonical” 3D density map of the molecule, and a parameterized latent space of deformation fields encoding non-rigid motion. The deformation fields “bend” the canonical density via convection, yielding all conformations captured by the model. In 3DFlex, the canonical density, the deformation field generator, and the latent coordinates of each particle image are jointly learned from the image data using a specialized training algorithm, without any prior knowledge about the flexibility of the molecule.

Our results on experimental cryo-EM data show that 3DFlex addresses the challenges of flexible refinement. We show that the model can jointly learn the structure of a flexible molecule, the underlying non-linear, non-rigid motion that unites its conformational landscape, and the positions of each single particle image on that landscape. Given a dataset of tri-snRNP spliceosome particles [24], 3DFlex learns non-rigid motions that range from slight deformations of fractions of an Angstrom in some regions, to the large motions of sub-units bending across a span of more than 20 Å. In doing so, the algorithm aggregates structural information from all conformations into a single, optimized density map that resolves high-resolution details in *α*-helices and *β*-sheets even in the flexible domains. In fact, 3DFlex can model continuous motion with enough precision to improve the resolution of small flexible parts that are otherwise poorly resolved in both conventional and local focused refinements. We demonstrate this ability with a dataset of TRPV1 ion-channel particles [10], where 3DFlex improves resolution and map quality of peripheral *α*-helices in the flexible soluble domains.

3DFlex is able to uncover motion in protein molecules, and to recover structural features that are usually lost in cryo-EM reconstructions of flexible specimens. This advance opens up many avenues of inquiry into further development of such methods, and into the study of biological mechanisms and function involving motion, at the frontier of both cryo-EM and structural biology generally.

## 2 3DFlex Model

3D Flexible Refinement (3DFlex) is a generative neural network method that determines the structure and motion of flexible protein molecules from cryo-EM images. Central to 3DFlex is the assumption that conformations of a dynamic protein are related to each other through deformation of a single 3D structure. Specifically, a flexible molecule is represented in terms of *i)* a **canonical** 3D map, *ii)* **latent coordinate** vectors that specify positions over the protein’s conformational landscape, and *iii)* a **flow generator** that converts a latent coordinate vector into a deformation field that convects the canonical map into the corresponding protein conformation. The canonical 3D map, the parameters of the flow generator, and a latent coordinate vector for each particle image are all model parameters that are initially unknown. They are jointly learned from experimental data.

Under the 3DFlex model (Figure 1), a single particle 2D image *I_i_* is generated as follows. First, the *K*-dimensional latent coordinates **z**_*i*_ of the particle are input to the flow generator *f_θ_*(**z**_*i*_). The generator provides a 3D deformation field, denoted **u**_*i*_(**x**), where **x** is a 3D position and *θ* denotes the parameters of the generator. The deformation vector field and the canonical 3D density map *V* are input to a convection operator, denoted *D*(**u**_*i*_, *V*), which outputs a convected density, denoted *W_i_*. The 2D particle image *I_i_* is then a CTF-corrupted projection of *W_i_*, plus additive noise *η*; i.e.,

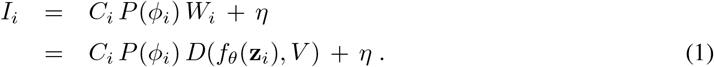

**Figure 1:**
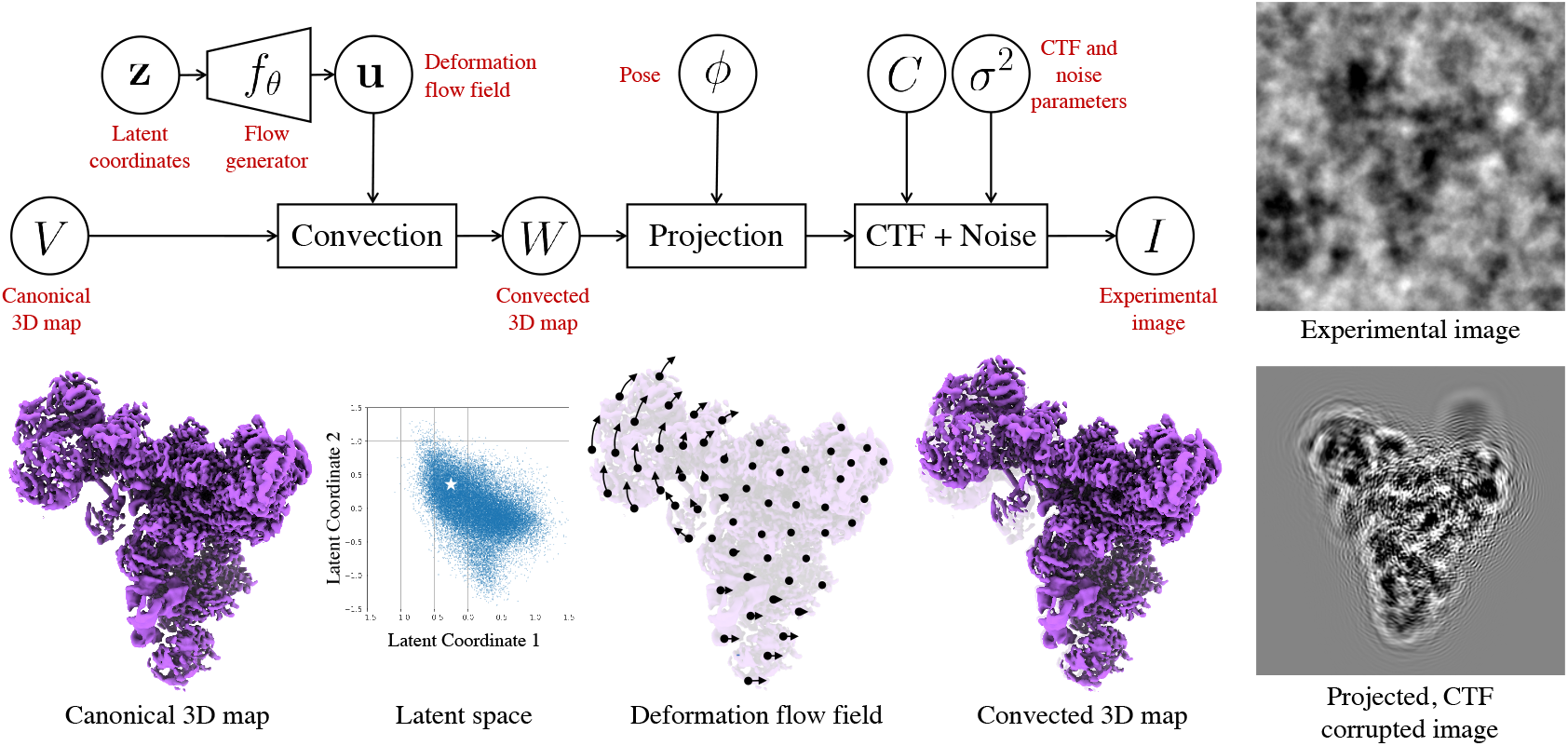
The 3DFlex model represents the flexible 3D structure of a protein as deformations of a single **canonical** 3D density map *V*. Under the model, a single particle image is associated with a low-dimensional **latent coordinate z** that encodes the conformation for the particle in the image. A neural **flow generator network***f_θ_* converts the latent coordinate into the flow field **u** and a convection operator then deforms the canonical density to generate a **convected** map *W*. This map can then be projected along the particle viewing direction determined by the pose *ϕ*, CTF-corrupted, and compared against the experimental image.

Here, *C_i_* denotes the CTF operator and *P* (*ϕ_i_*) is the projection operator for pose *ϕ_i_*, specifying the transformation between the microscope coordinate frame and the coordinate frame of the canonical map.

Fitting 3DFlex to experimental data entails optimizing the flow generator parameters *θ*, the canonical density map *V*, and the latent coordinates **z**_*i*_, in order to maximize the likelihood of the experimental data under the probabilistic model (Eq. 1). This is equivalent to minimizing the negative log-likelihood,

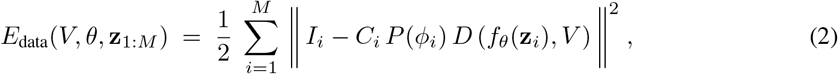

where *M* is the number of particle images. For notational simplicity we have assumed additive white noise, however it is straightforward to extent the formulation to handle colored noise, which is important in practice. For the development of 3DFlex below, we also assume poses *ϕ_i_* and CTF estimates are known, for example from a standard cryo-EM refinement algorithm, though these parameters could also be re-optimized in the 3DFlex model.

Within the 3DFlex formulation there are several important design choices that define the architecture of the 3DFlex model. Computationally determining structure and motion from noisy cryo-EM data is a challenging problem. As such, discussion of the design choices below provides insight into the working model, reflecting our exploration of different designs during the development of 3DFlex.

### Flow generator

We use a fully-connected deep neural network (a.k.a., a multi-layer perceptron, or MLP) with rectified linear activation functions (ReLUs) for the flow generator. The input **z** is the low-dimensional latent coordinate vector for a given image, and the output is a 3D flow field **u**(**x**). The number of hidden units per layer (typically 32-128) and the number of layers (typically 2-8) are adjustable hyperparameters. The final layer is linear (without biases or nonlinear activation).

The neural flow generator gives 3DFlex the capacity to learn complex, nonlinear deformation fields from data, and the inherent inductive bias of the architecture helps avoid over-fitting. Nevertheless, the data are noisy and the number of network parameters is large. Explicit regularization therefore plays a key role in reducing the risk of over-fitting (see below).

### Auto-decoder

The latent space in 3DFlex represents the conformational landscape, as different latent positions correspond to different deformations of the canonical map. A key step in training the 3DFlex model is to infer the latent coordinates (a.k.a. embedding) for each input image. In probabilistic terms, given an image *I*, the goal is to infer the posterior distribution *p*(**z***| I*) over latent coordinates, where high probability coordinates are those for which the flow generator and canonical map explain the image well. Equivalently, these are the latent coordinates that minimize Eq. 2.

Determining the exact posterior distribution is intractable for problems like 3DFlex, so instead one must resort to approximate inference. One approach, commonly used in variational auto-encoders (VAE) [16], is so-called *amortized variational inference*, in which a single feed-forward neural network (the encoder) is used to approximate the posterior for any image. Given the input image, the encoder learns to output a mean and covariance over latent coordinates. In essence, the encoder must learn to “invert” the generative model, predicting the most likely latent coordinates by looking at the input image. This approach has been used by deep-learning based heterogeneity methods proposed thus far [44, 3, 43]. In the context of 3DFlex, the encoder would be trained jointly with the flow generator and the canonical map, to maximize the likelihood of the particle images. VAE architectures are usually stable to train and inference can be fast, requiring just a single pass through the encoder network. They also incorporate a prior over latent coordinates that helps to regularize the structure of the latent space to be smooth and compact.

Amortized inference methods have several disadvantages. Primarily, it can be difficult for the encoder network to accurately approximate the posterior defined by the generative model [5]. In the context of 3DFlex, an encoder network must learn how to look at a single noisy 2D image and invert CTF corruption, 3D-to-2D projection, and 3D deformation. Motion must be learned in tandem with the generative model; when the flow generator shifts a particular sub-unit up or down, the encoder must simultaneously learn the same motion and how it appears from every 2D viewing direction, in order to infer the position of the sub-unit in each image. These requirements are difficult to fulfill, especially given the high noise level in experimental images. Indeed, we found amortized inference with an encoder to be insufficiently precise to resolve high-resolution structure and motion.

In 3DFlex we instead adopt a so-called **auto-decoder** model, where we perform inference by optimizing a point estimate of the latent coordinates independently for each image, taking advantage of the structure of the generative model directly. Although substantially more computationally expensive than amortized inference with an encoder network, this direct inference is more precise. Learning of motion only needs to happen once in the flow generator and naturally incorporates information across images and viewing directions. These characteristics enable 3DFlex to capture structure and motion with sufficient detail to resolve flexible protein regions to higher resolution than previous methods.

3DFlex has an end-to-end differentiable generative model, and so one can compute gradients of the data likelihood with respect to the latent coordinates for each image, and then use gradient-based optimization to perform inference. When the dimensionality *K* of the latent space is small enough, it is also possible to use coordinate-descent. We found the latter approach to be simpler and equally effective in our experiments.

### Noise injection and prior on latents

One benefit of explicitly modeling the posterior distribution *p*(**z***| I*), rather than optimizing a point estimate for **z** given *I*, is that accounting for uncertainty in **z** can regularize the model and encourage smoothness of the latent space by ensuring that nearby latent coordinates yield similar convected density maps after deformation.

In 3DFlex, we find that the heuristic method of directly adding noise to the point estimate during inference produces a similar positive effect. This method can be likened to variational inference with a Gaussian variational family with a fixed covariance, and has been used to regularize deterministic auto-encoders [11]. Finally, in addition to noise injection, we use a Gaussian prior on latent coordinates with unit variance to help control the spread of the latent embedding for a given dataset, and to center it at the origin in the latent space.

### Real-space vs Fourier-space

Algorithms for single-particle reconstruction commonly represent 3D maps and 2D images in the Fourier domain. Working in the Fourier domain reduces the computational cost of CTF modulation and image projection (via the Fourier-slice theorem). It also allows maximum-likelihood 3D reconstruction with known poses in closed-form (e.g., see [13, 35]). On the other hand, the convection of density between conformations is more naturally formulated as a real-space operation. Features and structures in the canonical density map *V* need to be shifted, rotated, and potentially deformed to produce densities consistent with the observed particles.

In 3DFlex we represent the canonical density *V* in real-space, as a voxel array of size *N*^3^. Convection and projection are performed in real-space, and in practice are combined into a single operator that does not store *W_i_* explicitly. Once the projected image of the convected map is formed, it is transformed to Fourier-space and CTF modulated, and transformed back to real-space to be used with the observed image for likelihood computation. Computationally, real-space convection and projection are far more expensive than Fourier-space slicing, and the FFT for CTF modulation must be applied for every image in the forward pass, and also in the backwards pass for computing gradients. Nevertheless we find that in 3DFlex, high-resolution 3D reconstruction of the canonical map is possible in real-space when using suitable optimization techniques (see below).

### Convection operator

Convection of density is an essential element of 3DFlex, modeling the physical nature of protein motion, thereby allowing high-resolution structural detail from experimental data to backpropagate through the model. There are several ways to construct a convection operator. One way is to express the flow field as a mapping from convected coordinates (i.e., of voxels in *W_i_*) to canonical coordinates. Convection then requires interpolating the canonical density *V* at positions specified by the flow field. However, in order to maintain conservation of mass the interpolated density must be modulated by the determinant of the Jacobian of the mapping, which can be challenging to compute and differentiate.

Instead, the flow in 3DFlex, **u**_*i*_(**x**), represents a forward mapping from canonical coordinates in *V* to the deformed coordinates in *W_i_*. This method naturally conserves density, as every voxel in *V* has a destination in *W_i_* where its contribution is accumulated through an interpolant function. The convected density at **x** can be written as

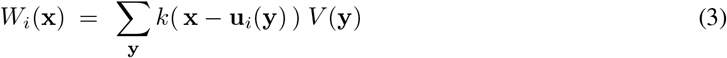

where **u**_*i*_ = *f_θ_*(**z**_*i*_), and *k*(**x**) is an interpolation kernel with finite support, and **y** is a spatial position that ranges over 3D voxels for the summation. In this case, divergence and convergence of the flow field must be treated carefully to avoid undesirable artifacts such as holes, Moiré patterns, and discontinuities. We found it critical to use high-order (eg. tricubic) interpolation and strong regularization (see below) to ensure accurate interpolation and artefact-free gradients.

### Regularization

As one adds capacity to a model like 3DFlex the propensity for over-fitting becomes problematic without well designed regularization. In early formulations of 3DFlex, over-fitting resulted in the formation of localized, high-density points (“blips”) in the canonical map, along with flow fields that translated these aberrations by large distances to explain noise in the experimental images. This problem was especially pronounced with smaller proteins, higher levels of image noise, and membrane proteins containing disordered micelle or nanodisc regions (i.e., structured noise). Over-fitting also occurs when the regularization is not strong enough to force the model to separate structure from motion. For example, rather than improve the canonical density with structure common to all conformations, the model sometimes learned to deform a low-resolution canonical density to create high-resolution structure (with highly variable local deformations).

To address these issues, 3DFlex exploits prior knowledge of smoothness and local rigidity in the deformation field. In particular, it is unlikely that natural deformations would involve large discontinuities in regions of high density; e.g. an *α*-helix should not be sheared into disjoint pieces. It is also unlikely that deformations will be highly non-rigid at fine scales in regions of high density; at the extreme, bond lengths should not stretch or compress substantially. With these intuitions we tried simple regularizers acting on flow fields defined at each voxel, like limiting the frequency content of the flow field, or penalizing its curvature. However these regularizers were difficult to tune and did not prevent over-fitting reliably.

3DFlex instead models flow generation using finite-element methods (FEM). A tetrahedral mesh covering regions of high density is generated in the canonical frame, based on a preliminary consensus refinement. The deformation field is parameterized by a 3D flow vector at each vertex of the tetrahedral mesh. The deformation field is then interpolated using linear FEM shape functions within each mesh element. Smoothness is a function of the size of mesh elements (an adjustable parameter) and is enforced implicitly through interpolation and the fact that adjacent elements share vertices.

We also encourage local rigidity of the flow in each mesh element. The deformation field within the *j*th tetrahedral element for image *i*, denoted **u**_*ij*_ (**x**) can be written as a linear mapping:

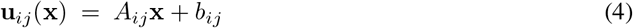

where matrix *A* and vector *b* are uniquely determined from 3D flow vectors at the element vertices. We quantify local non-rigidity in terms of the distance between *A* and the nearest orthogonal matrix (in a mean squared-error sense [2, 14]). In particular, we measure the squared deviation of the singular values of *A* from unity. Letting 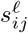 be the *ℓ*th singular value of *A_ij_*, we express the local rigidity regularization loss as

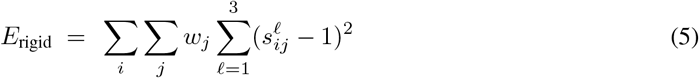

where *w_j_* are weights defining the strength of the prior within each mesh element, based on the density present within the *j*th mesh element. The densest elements have weight one and empty elements have weight zero. This weighting ensures that deformation fields are permitted to compress and expand empty space around the protein.

### Optimization of flow and structure

3DFlex is end-to-end differentiable, allowing gradient-based optimization to be used to train the flow generator and learn the canonical density that best explains the data. We use either Adam [15] or Stochastic Gradient Descent (SGD) with Nesterov acceleration [41] with minibatches of size at least 500 due to the high levels of image noise. Inference of the latent coordinates for each image in a minibatch is performed prior to computing gradients with respect to the the canonical density and flow parameters. The loss function is a weighted sum of the data log likelihood (Eq. 2) and the non-rigidity penalty (Eq. 5):

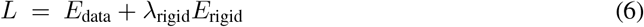

During optimization, we use frequency marching, learning the model in a coarse-to-fine manner. The canonical density *V* is constrained to be low-pass, with an upper frequency band-limit that increases over iterations. The frequency and learning rate schedule, and λ_rigid_, must be tuned for each dataset in our current implementation. Optimization is done with a box size, *N* = *N_L_*, that is typically smaller than the raw size of the particle images *N_H_*. As such, optimization only uses information below the Nyquist frequency for the smaller box size. During optimization, it is also possible to apply a real-space mask to the flow generator output to ensure that the deformation is zero except in a region of interest, e.g. to exclude motion of a micelle or nanodisc.

To initialize training we set the canonical density *V* to be a low-pass filtered version of a conventional consensus refinement map given the same particle images. The parameters of the flow generator are randomly initialized. The latent coordinates for the particles are either initialized to zero or to the output of another heterogeneity embedding method. In some experiments, especially on smaller, low SNR particles, we find that initializing with latent coordinates from 3D Variability Analysis (3DVA) [28] in cryoSPARC [29] improves results.

We also find that simultaneously training the canonical density *V* and flow generator parameters *θ* leads to over-fitting after thousands of gradient iterations, despite strong regularization. In these cases, we opt to initially train *V* and *θ* with latent coordinates *z_i_* fixed to their initial values from 3DVA for 5 epochs. Then, we unlock the latent coordinates and perform latent inference for each minibatch while updating *θ*, but with *V* fixed, for 10 epochs. Then, we alternate to updating *V* with *θ* fixed for 10 epochs, and repeat until convergence.

### High resolution refinement and validation

With the ability of 3DFlex to capture motion and latent coordinates of each particle image, it becomes possible in principle to attempt recovery of high-resolution detail in flexible parts of protein molecules that would otherwise be blurred in standard reconstructions.

3DFlex is optimized at a small box size, *N* = *N_L_*. Once optimization has converged, we freeze the flow generator parameters *θ* and the latent coordinates 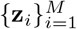, and then transfer them to a new model at full resolution, with *N* = *N_H_*. We partition the particles using the same split that was used in the consensus refinement (from which we obtained the poses *{ϕ_i_}*). For each half-set we initialize the canonical density *V* to zero, and re-optimize it at full box size *N_H_*, with the other parts of the model fixed. In the same way as established reconstruction validation methods [12, 38], the two resulting half-maps can be compared via FSC. Correlation beyond the training-time Nyquist resolution limit indicates that true signal was recovered in common from both separate particle sets, as opposed to spurious correlation or over-fitting of the model. This correlation serves to validate the improvement in reconstruction of flexible regions.

To this end, we need to optimize *V* at high resolution under the full 3DFlex model for each half-set. We initially experimented with localized Fourier-space reconstructions but encountered issues with non-rigidity and curvature. Perhaps unsurprisingly, we also found that minibatch SGD methods for directly optimizing *V* did not yield high quality results, potentially because noise in minibatch gradient estimation is problematic for this task. Instead, we were able to solve the problem using full-batch L-BFGS [25] in real-space. This approach is substantially more expensive than Fourier-space reconstruction, and requires many iterative passes over the whole dataset. However, it is notable in that it allows 3DFlex to solve high-resolution detail in all flexible parts of the protein simultaneously, without making assumptions of local rigidity or smoothness.

## 3 Experimental Results

We consider the application of 3DFlex to two experimental cryo-EM datasets, demonstrating the ability of the method to resolve multiple dimensions of non-rigid protein motion with sufficient fidelity to improve reconstruction resolution of flexible parts. The first dataset contains tri-snRNP spliceosome particle images 4, and the second contains TRPV1 ion-channel particle images [10].

For each dataset, we first compute a rigid consensus refinement using all particle images. Non-uniform refinement [30] is used to improve image alignments and overall resolution. The resulting particle images and poses *ϕ_i_* are fixed during training of the 3DFlex model. No prior information is provided about the type or form of heterogeneity in each dataset. 3DFlex is run with a real-space mask that excludes solvent in the canonical density *V*. For membrane proteins a separate mask is used to enforce zero deformation in the region of detergent micelle or lipid nanodisc. For TRPV1, 3D Variability Analysis [28] was first run on the particle images to generate initializations for latent coordinates *z_i_*, again without any prior information about heterogeneity. Each experiment was run on a single NVIDIA V100 GPU.

For visualization of canonical density, half-maps from 3DFlex were combined, filtered by their FSC curve, and B-factor sharpened. To display conformational changes in figures, we selected points in the 3DFlex latent space, e.g., *z*_display_, and then generated the corresponding convected densities, *W*_display_ = *D*(*f_θ_*(*z*_display_)*, V*). These densities are rendered overlaid in multiple colors, and with reference position guide markers help visualize the motion. Videos depicting continuous motion and non-rigid deformation in much greater clarity are available as Supplementary Videos and also at https://cryosparc.com/3dflex.

### 3.1 snRNP spliceosome: Large non-rigid deformations of a complex

The U4/U6.U5 tri-snRNP complex represents a large part of the spliceosome. It has several moving parts, linkages, and flexible domains [24]. We process a dataset of 138,899 snRNP particles (EMPIAR-10073). The raw particle images have a box size of 380 pixels, and a pixel size of 1.4Å and are first processed through heterogeneous refinement in *cryoSPARC* to separate broken particles that are missing the “head” region. This yields 102,500 final particles. They are downsampled to a box size of 180 pixels (pixel size 2.95Å) and used to train 3DFlex. Given the optimized latent coordinates and trained flow generator, the original raw particle images, at full resolution (380 pixels), are then used to reconstruct the canonical density (in two separate half-maps) to high resolution using L-BFGS.

3DFlex training began with random initialization of the latent coordinates and the flow generator parameters. We experimented with several flow generator architectures, and latent dimensionality between *K* = 1 and 5. Larger latent spaces allowed 3DFlex to discover more distinct motions of the complex, and we show results from a *K* = 5-dimensional latent space, with a 6-layer MLP with 64 units in each hidden layer and output layer for the flow generator network. A tetrahedral mesh with 1601 vertices and 5859 cells was created to cover the density. The flow generator outputs 3D displacement vectors at these vertices.

3DFlex recovers five dimensions of latent motion (Figure 2, Supplementary Video 1), with each dimension explaining a different type of bending or twisting in the molecule. There are two large moving parts, namely, the head region and foot region. Both are attached to the more rigid central body. In the learned deformation fields from 3DFlex, the foot region largely moves as a rigid sub-part, with a hinge-like linkage to the body. The head region has significant internal flexibility and a large range of motion. All five latent directions encode some motion in the head region. Notably, 3DFlex recovers the motion of all regions from a random initialization without any labels defining parts or linkages.

**Figure 2:**
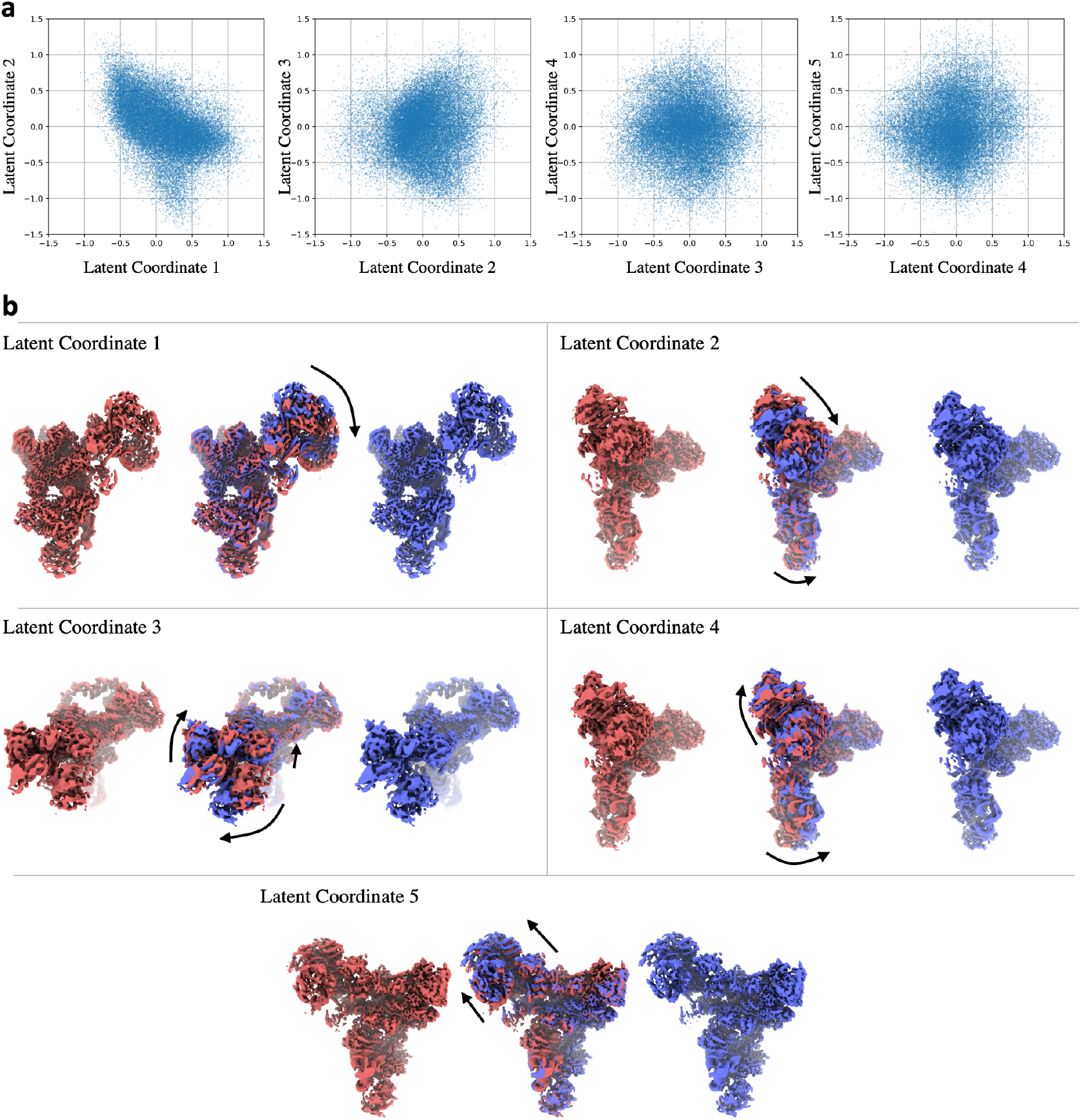
Results of 3DFlex with a *K* = 5-dimensional latent space on 102,500 particle images of an snRNP Spliceosome complex [24], demonstrating the capacity for 3DFlex to resolve multiple modes of non-rigid deformation simultaneously. (Supplementary Video 1, also at https://cryosparc.com/3dflex, depicts motion with much greater clarity) **a**: Scatter plots showing the final distribution of particle latent coordinates across the dataset. **b**: Convected densities from 3DFlex at minus one (red) and plus one (blue) standard deviations in the latent space, along each of the five latent dimensions. Each dimension resolves a different type of motion within the same model.

In addition to the 3D motion between conformations and the canonical map, 3DFlex also recovers high-resolution detail in the canonical map (Figure 3, Supplementary Video 1). Individual *α*-helices can be seen translating several Angstroms while retaining side-chain features. Likewise, a *β*-sheet in the flexible head region is resolved with separated *β*-strands, despite the presence of non-rigid motion. 3DFlex only has access to experimental data at a maximum Nyquist resolution of 5.9Å during training of the 3DFlex model with the flow generator, and so these structural features represent additional signal that is resolved from the data due to the accuracy of the recovered motion.

**Figure 3:**
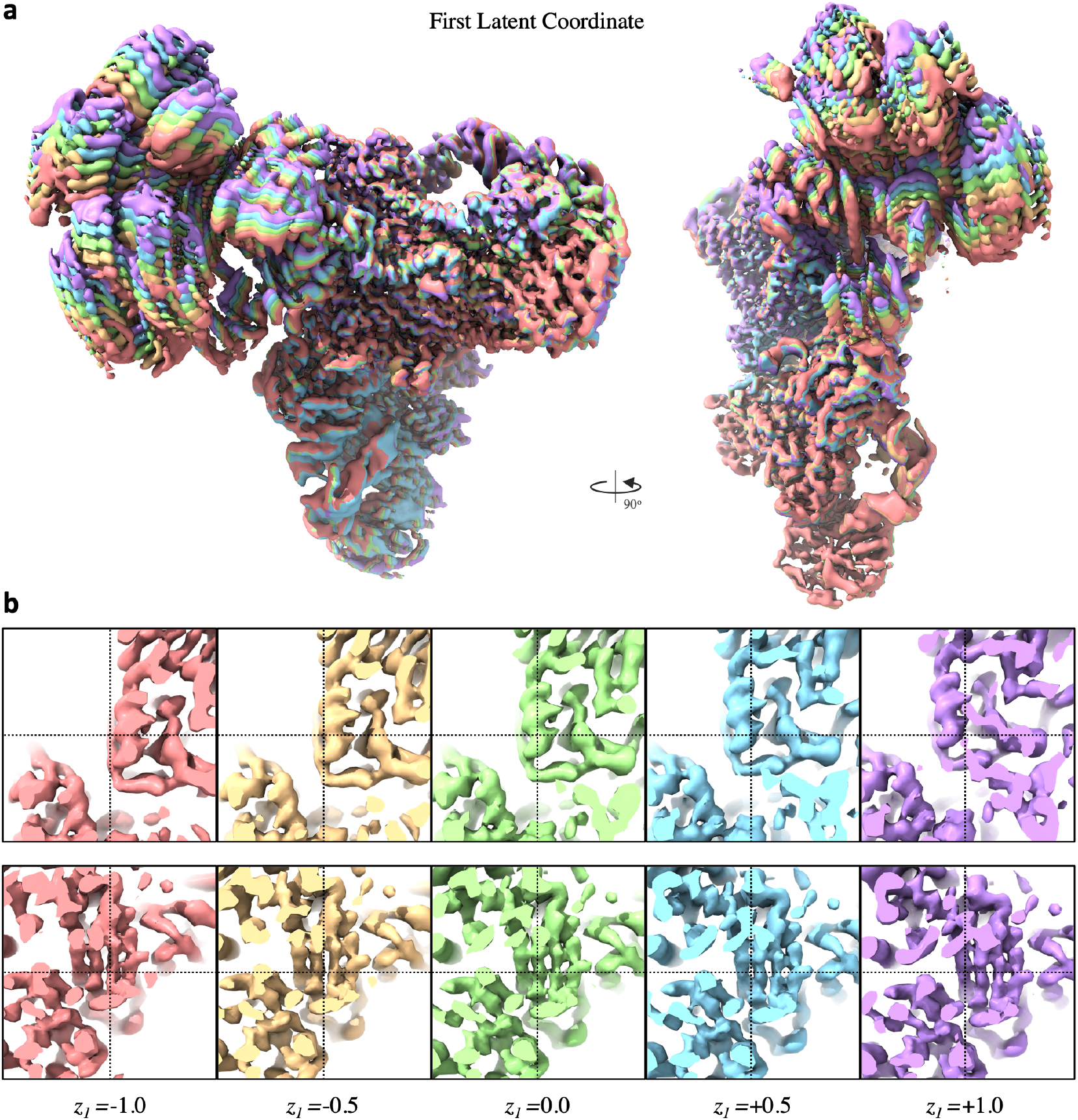
Results of 3DFlex on 102,500 particle images of an snRNP Spliceosome complex [24], demonstrating the capacity for 3DFlex to resolve detailed non-rigid motion and high-resolution structure simultaneously. (Supplementary Video 1, also at https://cryosparc.com/3dflex, depicts motion with much greater clarity) **a**: Series of convected densities from the 3DFlex model, at latent coordinates along the first latent dimension. **b**: Same as **a** but with focus on key structural details. Top row: An *α*-helix in the head region of the protein that translated by several Angstroms. Bottom row: A *β*-sheet in the head region that translates and deforms.

As expected, in regions of substantial motion and flexibility, differences between a static conventional refinement and 3DFlex are dramatic (Figure 4). For example, local resolution in the center of the head region is improved from 5.7Å to 3.8Å. For a complex as large as the snRNP, it is worth noting that one could create manual masks around regions that are expected to be rigid, and then perform local or multi-body refinement [37]. Such refinement techniques can improve resolution and map quality in domains like the foot region, which remains rigid despite motion relative to the remainder of the molecule. In contrast, 3DFlex does not require manual masking or prior knowledge about the motion of the molecule. It can detect and then correct for non-rigid flexibility across the entire molecule at once, including the head region, which is considerably less rigid than the foot.

**Figure 4:**
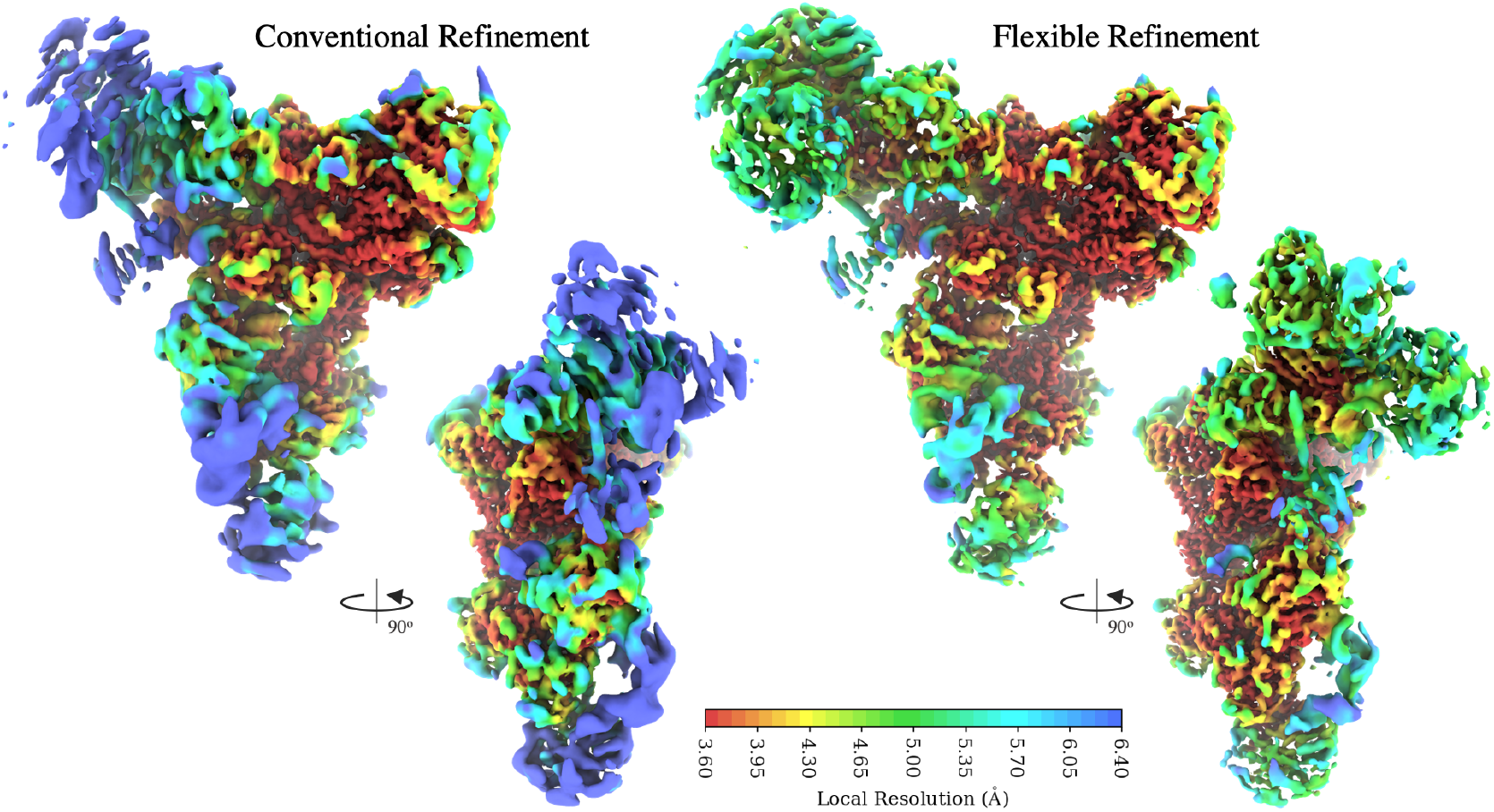
Results of 3DFlex with a *K* = 5-dimensional latent space on 102,500 particle images of an snRNP Spliceosome complex [24]. Left: density map from conventional refinement colored by local resolution. Right: canonical density map from 3DFlex, colored on the same local resolution color scale. The two maps are filtered by local resolution to aid in visualizing weak density in low-resolution areas in the conventional refinement.

### 3.2 TRPV1 ion channel: Capturing flexible motion improves resolution

The TRPV1 ion channel is a 380 kDa tetrameric membrane protein that acts as a heat- and capsaicin-activated sensory cation channel [10]. We process a dataset of 200,000 particle images of TRPV1 in nanodisc (EMPIAR-10059) with a box size of 224 pixels and pixel size of 1.21Å. These particles are downsampled to 128 pixels (pixel size 2.15Å) and used to train 3DFlex. The original raw particle images are then used to reconstruct the canonical density (in two separate half-maps) to high resolution, once 3DFlex has learned the motion.

We first run 3D Variability Analysis [28] with *K* = 2 components to provide initialization for latent coordinates **z**_*i*_ for 3DFlex. The flow generator, a 3-layer MLP with 32 units in each hidden and output layer, is randomly initialized. 3DFlex is trained for 5 epochs with the latent coordinates fixed. The latent coordinates are then unlocked and are optimized at each iteration. Training proceeds for 10 epochs at a time, alternating between fixing the canonical density and fixing the flow generator, for a total of 50 epochs.

The final result is a 3DFlex model that has captured *K* = 2 types of flexible coordinated motion amongst the four peripheral soluble domains of the ion channel (Figure 5, Supplementary Video 2). Along the first latent dimension, each pair of opposing subunits bends towards each other while the other pair bends apart. The second motion involves all four subunits twisting concentrically around the channel’s pore axis. In both cases, the peripheral-most helices move by approximately 6Å. Both motions are non-rigid and involve flexure of substantial regions of the protein density.

**Figure 5:**
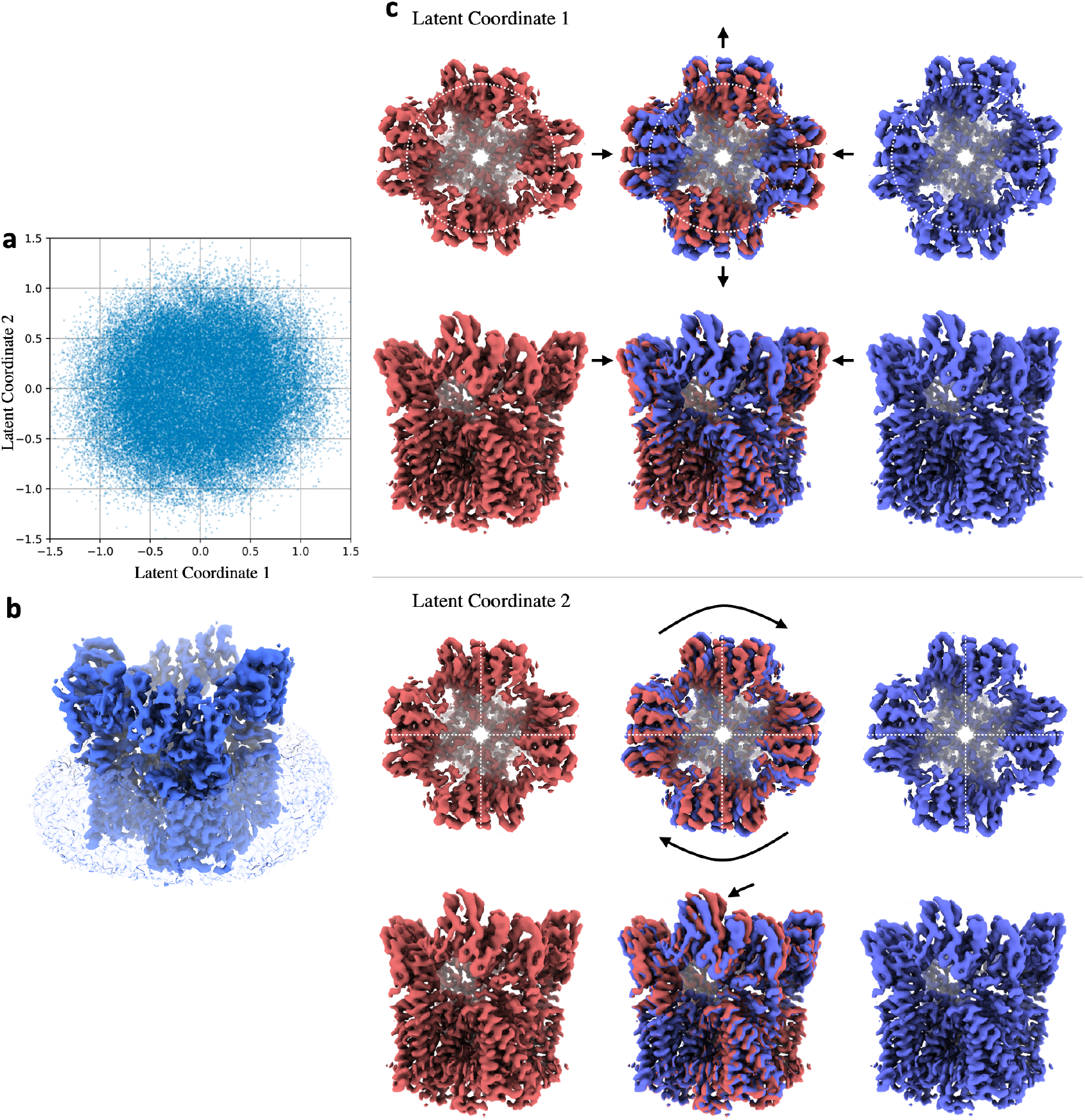
Results of 3DFlex with a *K* = 2-dimensional latent space on 200,000 particle images of a TRPV1 ion channel protein [10], demonstrating the capacity for 3DFlex to resolve detailed motion of smaller, membrane proteins. (Supplementary Video 2, also at https://cryosparc.com/3dflex, depicts motion with much greater clarity) **a**: Scatter plots showing the final distribution of particle latent coordinates across the dataset. **b**: Canonical density that is determined by 3DFlex. The micelle, shown in transparent blue, is not excluded in the density but is masked to have zero deformation, so that 3DFlex focuses on motion of the protein density. **c**: Convected densities from 3DFlex at minus one (red) and plus one (blue) standard deviations in the latent space, along each of the two latent dimensions. The first dimension reveals a motion where opposite soluble domains move together or apart. The second dimension reveals a motion where all four soluble domains twist around the axis of the central pore.

In a conventional refinement of the TRPV1 channel, these motions are detrimental to reconstruction quality and resolution (Figure 6, Supplementary Video 3). Several *α*-helices in the soluble region are so poorely resolved that helical pitch is barely visible. Local resolution reaches 2.8Å in the rigid core of the channel, but only 4Å at the periphery. 3DFlex, on the other hand, estimates and accounts for the motion of these domains, and substantially improves resolution and map quality. 3DFlex only has access to experimental data up to a maximum Nyquist resolution of 4.3Å during training, but FSC and local resolution measurements using the two separate half-set reconstruction in 3DFlex show that it recovers consistent structural information beyond this resolution. Local resolutions in peripheral helices improve to 3.2Å revealing helical pitch and side chain details.

**Figure 6:**
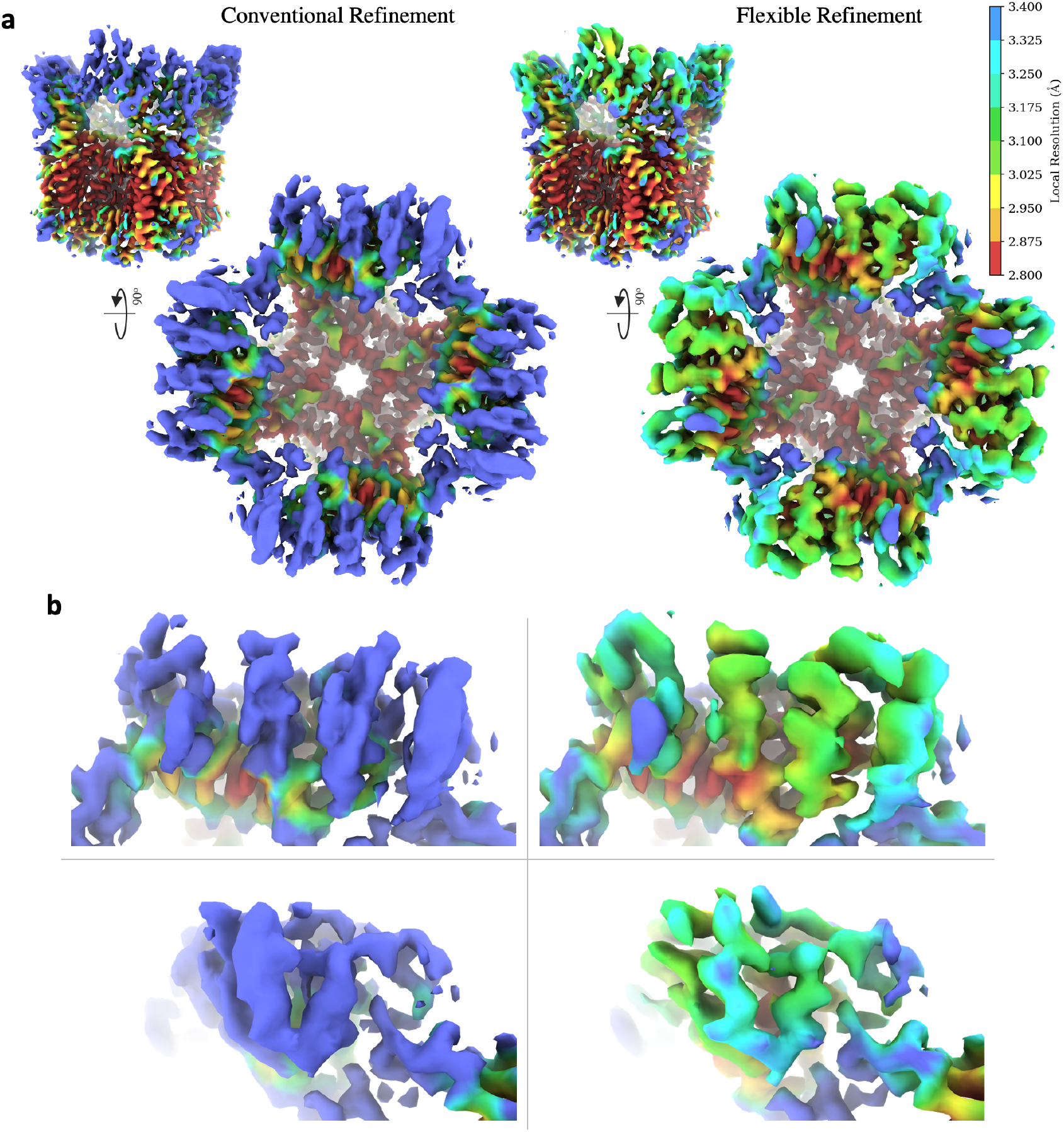
Results of 3DFlex with a *K* = 2-dimensional latent space on 200,000 particle images of a TRPV1 ion channel protein [10]. (Supplementary Video 3, also at https://cryosparc.com/3dflex, depicts the comparison with much greater clarity) **a**: Left: density map from conventional refinement colored by local resolution. Right: canonical density map from 3DFlex, colored on the same local resolution color scale. The two maps are filtered and sharpened identically and displayed at the same threshold level, so that visual comparison of map quality is possible. The 3DFlex result shows clear improvement in map quality and local resolution in peripheral flexible domains of the protein. **b**: Detail views showing improvement in helical density in the flexible soluble domains.

The two separate half-set reconstructions from 3DFlex allow us to use established validation procedures to measure the improvement that derives from the motion estimation. Figure 7a shows that the global FSC curve of the entire density improves slightly in 3DFlex compared to conventional refinement. This difference indicates that in the rigid core region of the molecule, 3DFlex has not lost any structural information. To investigate the effect in the peripheral domains, we constructed a soft-edged mask around one of the flexible domains (Figure 7c) and tested the mask for tightness using noise-substitution [33]. Computing FSC curves within this mask (Figure 7) shows that 3DFlex improves the average resolution from 3.4Å to 3.2Å as well as increasing the SNR at low and medium resolutions. This improvement means that 3DFlex has resolved more structural information than conventional refinement for this flexible protein, and validates that the motion learned by 3DFlex is a better model of the particle than a null hypothesis of no motion.

**Figure 7:**
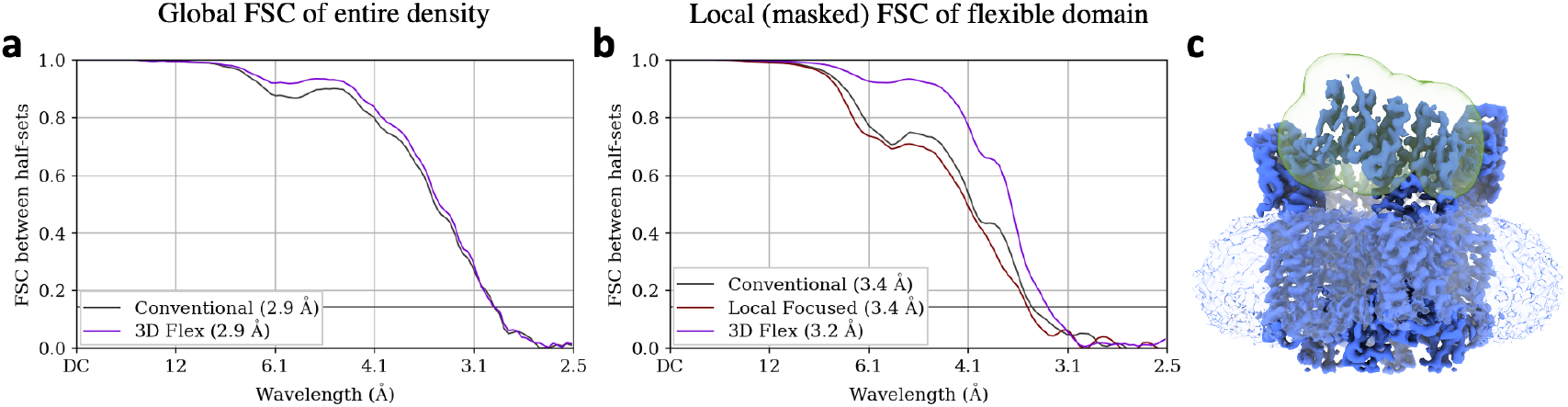
Validation and resolution estimation results of 3DFlex with a *K* = 2-dimensional latent space on 200,000 particle images of a TRPV1 ion channel [10]. FSC curves are measured between half-sets of particles that are used to compute two half-map reconstructions. 3DFlex only has access to experimental data up to a Nyquist resolution limit of 4.3Å during training. Therefore, any correlation beyond this resolution in half-set reconstructions indicates resolved signal rather than spurious correlation or over-fitting. **a**: FSC of the entire density of the ion channel, showing that 3DFlex resolves high-resolution details in the rigid core of the protein. **b**: FSC using a mask around one of the flexible peripheral domains in the soluble region of the ion channel. The mask is depicted in **c** and is soft-edged. In this region, 3DFlex provides a substantially improved FSC curve and resolution estimate of 3.2Å versus 3.4Å for conventional refinement. Notably, a local focused refinement using the same mask (**b**, red) is unable to improve resolution beyond the conventional refinement result due to the small size and flexibility of the region.

3DFlex improves the reconstruction of this flexible protein by explicitly modelling non-rigid deformation. As a baseline, we also perform a local focused refinement using the same mask (Figure 7c) to isolate one soluble domain. Local refinement is unable to improve the density or resolution of the domain beyond conventional refinement (Figure 7b). This result is expected, as each soluble domain is less than 50 kDa in size and deforms flexibly. We believe that this comparison illustrates an additional advantage of 3DFlex. Unlike local and multi-body refinement methods that assume rigidity and attempt to fit separate pose parameters for each masked region, 3DFlex exploits correlations between different moving parts that move together, making it possible to infer the position of all parts, even though individually each is too small to align reliably. In the case of TRPV1, the four soluble domains deform in different directions by different amounts, but 3DFlex infers their positions in a given image jointly.

## 4 Discussion

*3D Flexible Refinement* (3DFlex) can recover detailed non-rigid protein motion from single particle cryo-EM data of both large complexes and smaller membrane proteins. At the same time, the learned motion is used to improve resolution and map quality of flexible regions in a single combined canonical density map that aggregates structural information from across a protein’s conformational landscape. The learned motion model and associated distribution of latent coordinates of individual particle images may shed light on biological function and protein dynamics.

### Related methods for single-particle cryo-EM heterogeneity

The most common methods currently used to resolve heterogeneity in cryo-EM data are based on discrete 3D classification [12, 34, 36, 39]. Discrete models approximate continuous heterogeneity by quantizing the landscape and grouping together particles that are assumed to have the same conformation. As such, within a single 3D class there may be substantial conformational variability. Perhaps more importantly, 3D classification methods do not aggregate structural information across discrete classes, nor do they model or exploit motion. Advanced methods [43] can provide finer and more elaborate discrete clustering with relatively large numbers of classes, but still do not aggregate structural information, and thus require large datasets to partition into many classes. In contrast, 3DFlex is highly data-efficient, as each particle contributes to the canonical density regardless of its conformational state.

Local, focused refinement [12, 36] and multi-body refinement [23] methods allow high-resolution refinement of flexible molecules by assuming that the molecule is composed of a small number of rigid parts. These parts must be identified and separated by manually creating masks for each region. And each part must have sufficient molecular weight (typically 150kDa or more) for accurate individual alignment independent of the rest of the structure [37]. Non-rigidity within each part is not modelled. 3DFlex does not require the manual creation of masks and can recover motion and structure of non-rigidly deformable parts across an entire molecule at once, although this recovery depends on mesh granularity, one of the model hyper-parameters.

Techniques like normal-modes analysis [6] make assumptions about the local energy landscape and dynamics of a protein around a base state of a molecule to predict flexibility. Methods have been proposed to exploit these fixed prior models to recover improved density maps from cryo-EM data of flexible molecules [40]. In contrast, 3DFlex does not presuppose knowledge of the energy landscape, bending, or flexing of the molecule. Rather, it learns this information from the image data.

Several promising density-based models have been developed to learn continuous heterogeneity from cryo-EM images. The simplest of these, closely related to normal modes analysis, are methods based on eigen-analysis [1, 27, 28, 42] that model conformational landscapes as linear subspaces in the space of 3D density. More advanced techniques use non-linear manifold embedding [7, 8, 9, 20, 22] or deep generative models [44, 43] to construct a non-linear manifold in the space of 3D density. These methods do succeed in modelling general continuous heterogeneity (as opposed to discrete or rigid approximations) but do not have a notion of protein motion or the preservation of local geometry. Instead, density-based models capture conformational change by adding and subtracting density in different areas of a 3D structure. Unlike 3DFlex, they have no inherent representation of convection. As such, structural information is not aggregated across the conformational landscape and reconstruction quality is not improved.

Finally, methods have also been proposed that can, in principle, model heterogeneity using an underlying notion of protein motion. Hypermolecules [18] represent heterogeneous protein density in a higher-dimensional space that has the potential to capture motion-based deformation and structure together, although this has yet to be demonstrated on experimental data. Another recently proposed method fits a weighted sum of Gaussian densities to a rigid consensus reconstruction of particle images. Then, a deep generative model is used to adjust the positions and amplitudes of the individual Gaussian components in order to model continuous conformational change of protein density. Shifting the positions of Gaussian components can be thought of as convection, and in this way, the method is similar to 3DFlex. However, unlike 3DFlex, the Gaussian method uses an auto-encoder for training and does not directly encode particle images, but rather uses a perturbation-like representation of the images relative to the initially-learned neutral Gaussian positions. These architecture choices, and other limitations mentioned in [3], mean that while the method can model motion at coarse resolutions on experimental data, it does not improve reconstruction quality of the aggregate density beyond the quality of the initial rigid consensus reconstruction.

### Future work

The introduction of 3DFlex opens many avenues for further research. These directions include further methods development, as well as validation techniques. There are also opportunities for approaches to further analysis and interpretation of results, along with techniques to enable researchers to uncover biological insight.

3DFlex can be improved and extended in several ways. Although we have described several variations of architecture and optimization choices, there are many more to consider. For instance, it is not necessary to use a real-space voxel representation for the canonical density. Neural fields or implicit functions [4, 21, 26, 32] may serve as a natural alternative. They can also be directly optimized with gradient-based algorithms, but do not have the limitations of voxel representations in terms of spatially uniform resolution, sampling and interpolation. Similarly, alternative representations of flow fields, beyond per-voxel flow and tetrahedral mesh deformation, may be useful. Regularization of 3DFlex, both in terms of the flow generator and the canonical density, can make use of structurally-aware priors. The flow generator can also be expanded to allow for certain motions (e.g. rotary) to be more naturally encoded, and the architecture can be expanded to more gracefully handle compositional variability.

There are few validation techniques for existing continuous heterogeneity methods, but the ability of 3DFlex to resolve detailed motion will necessitate the development of methods capable of validating that the motion and deformation fields are in fact correct and supported by the image data.

The interpretation of 3DFlex results will also require further development. For instance, although tempting, it is unclear how one should relate the continuous probability distribution of particle images in the 3DFlex latent space to a physically meaningful notion of energy via a Boltzmann distribution. This is because the non-linear capacity of the flow generator means that relative distances and volumes (and therefore probability density) in the latent space are arbitrary.

Demonstrating the capacity of 3DFlex to learn motion and structure from cryo-EM data also raises questions about optimizing upstream methods in sample preparation and data collection to maximize the available information in particle images. For instance, 3DFlex may be able to yield significant biological insight when combined with time-resolved sample preparation or multiple datasets collected under different sample conditions or in the presence of different ligands.

## Supporting information

Supplementary Video 1 - snRNP motion

Supplementary Video 2 - TRPV1 motion

Supplementary Video 3 - TRPV1 reconstruction

## Code Availability

We plan to release an experimental version of 3DFlex in *cryoSPARC*. We also ask those interested in collaborating on analyzing datasets of flexible proteins to please contact us.

## Acknowledgements

We thank the entire team at Structura Biotechnology Inc. that designs, develops, and maintains the *cryoSPARC* software system on top of which this project was implemented and tested. Resources used in this research were provided, in part, by the Province of Ontario, the Government of Canada through NSERC and CIFAR, and companies sponsoring the Vector Institute.

